# An updated AL-Base reveals ranked enrichment of immunoglobulin light chain variable genes in AL amyloidosis

**DOI:** 10.1101/2024.09.11.612490

**Authors:** Gareth Morgan, Allison N. Nau, Sherry Wong, Brian H. Spencer, Yun Shen, Axin Hua, Matthew J. Bullard, Vaishali Sanchorawala, Tatiana Prokaeva

## Abstract

**Background:** Each monoclonal antibody light chain associated with AL amyloidosis has a unique sequence. Defining how these sequences lead to amyloid deposition could facilitate faster diagnosis and lead to new treatments.

**Methods:** Light chain sequences are collected in the Boston University AL-Base repository. Monoclonal sequences from AL amyloidosis, multiple myeloma and the healthy polyclonal immune repertoire were compared to identify differences in precursor gene use, mutation frequency and physicochemical properties.

**Results:** AL-Base now contains 2,193 monoclonal light chain sequences from plasma cell dyscrasias. Sixteen germline precursor genes were enriched in AL amyloidosis, relative to multiple myeloma and the polyclonal repertoire. Two genes, *IGKV1-16* and *IGLV1-36*, were infrequently observed but highly enriched in AL amyloidosis. The number of mutations varied widely between light chains. AL-associated κ light chains harbored significantly more mutations compared to multiple myeloma and polyclonal sequences, whereas AL-associated λ light chains had fewer mutations. Machine learning tools designed to predict amyloid propensity were less accurate for new sequences than their original training data.

**Conclusions:** Rarely-observed light chain variable genes may carry a high risk of AL amyloidosis. New approaches are needed to define sequence-associated risk factors for AL amyloidosis. AL-Base is a foundational resource for such studies.

## Introduction

Deposition of monoclonal immunoglobulin light chain (LC) proteins as amyloid fibrils leads to progressive toxicity and organ failure in individuals with AL amyloidosis [1]. LCs are secreted from an aberrant clonal population of B lymphocytes, most commonly bone marrow plasma cells, both bound to heavy chains as antibodies and as unbound free LCs. AL amyloidosis is one of a spectrum of conditions called plasma cell dyscrasias (PCD), which includes the more common blood cancer multiple myeloma (MM). The amino acid sequence of the LC influences the propensity of the protein to aggregate as well as the structure and interactions of the resulting amyloid fibrils [2,3]. Aggregation is concentration-dependent and rarely observed in the absence of a monoclonal LC. However, despite the generally higher levels of circulating monoclonal LC in MM, amyloid deposition is observed only in a subset of patients and may not lead to organ involvement [4,5]. The pathological consequences of each clone, referred to as amyloidogenicity, appear to be determined by both a LC’s concentration and physicochemical properties, which are defined by protein sequence [2,3]. Mechanistic understanding of the relationship between LC sequence and pathology could allow to identify the potentially dangerous LCs before patients become symptomatic, predict which organs may be at risk of involvement, and support the development of new therapies.

Each B cell expresses a single LC protein sequence derived from one of two germline loci, *IGK* or *IGL*, which encode κ or λ LCs, respectively. Within these loci, precursor variable (*IGV*_*L*_), joining (*IGJ*_*L*_) and constant (*IGC*_*L*_) gene regions are rearranged and subject to somatic hypermutation to create a functional *IGV*_*L*_-*IGJ*_*L*_-*IGC*_*L*_ light chain gene [2]. These processes lead to enormous sequence diversity, so that each patient has a unique monoclonal LC [6]. This sequence diversity is concentrated in the antigen-binding variable (V_L_) domain of the LC protein, residues from which form the structured core of AL amyloid fibrils [7]. In this manuscript, rearranged genes are referred to as *IGLV*-*IGLJ* or *IGKV*-*IGKJ* nucleotide sequences, which encode the V_L_-domains of κ and λ LC proteins, respectively. Subscripts denote light chain genes of either isotype (e.g., *IGV*_*L*_) to distinguish them from λ-encoding genes (e.g.*IGLV*). Protein sequences are referred to as κ and λ LCs, or as V_L_-domains. Light chain genes and the proteins derived from them are named and numbered according to the Immunogenetics (IMGT) nomenclature [8].

Numerous studies have sought LC sequence features such as gene us and specific mutations associated with AL amyloidosis [6,9–16]. Monoclonal *IGV*_*L*_-*IGJ*_*L*_-*IGC*_*L*_ gene and LC protein sequences can be determined by sequencing of mRNA extracted from plasma cells, or peptide sequencing using Edman degradation or mass spectrometry, respectively [16–18]. New approaches have adapted high throughput sequencing methods to significantly increase the volume of sequenced *IG*_*L*_ genes [19–21]. The clonal LC is encoded by abundantly expressed mRNA in bone marrow plasma cells, although some patients feature oligoclonal LCs [22]. The monoclonal LC protein encoded by these genes can be identified in tissue amyloid deposits, blood or urine [23,24], which allows sequences identified by different approaches to be studied together. Studies have observed an over-representation of LCs derived from a particular subset of precursor genes in patients with AL amyloidosis [6,12–16,25].

To facilitate efforts in identification of sequence markers of LC amyloid propensity, the AL-Base database and website were launched by the Boston University Amyloidosis Center in 2009 [6]. This publicly accessible resource collects nucleotide and protein sequences known to be associated with AL amyloidosis and other PCDs including MM, light chain deposition disease, and others. We now present an expanded and updated AL-Base, incorporating sequences reported in the literature since the previous update. These monoclonal sequences are analyzed together with a cohort of >8 million polyclonal LC sequences obtained from the Observed Antibody Space resource [26,27], revealing details of the relative enrichment of *IGV*_*L*_ genes in AL amyloidosis. Sequence-derived parameters that have been reported to differ between amyloidogenic and non-amyloidogenic LCs are also investigated.

## Methods

### AL-Base website

AL-Base has been completely rebuilt as a standalone web application using the Microsoft ASP.NET framework. It is available online at http://albase.bumc.bu.edu/aldb. Sequences are stored in a SQL Server database. Alignments to the closest germline gene were determined for each sequence as described below and are stored locally. Protein multiple sequence alignments of V_L_-domain can be created directly from these alignments. De novo multiple sequence alignments of protein or nucleotide sequences can be determined externally, using the ClustalW server at https://www.genome.jp/tools-bin/clustalw. A basic local alignment search tool (BLAST) database [28] allows homologous sequences to be identified.

### Classification of LC sequences

Each sequence was assigned to a clinical category and subcategory, based on the case description provided in the source paper or deposition. Categories define whether the sequence originated from a PCD and if so, whether amyloid was present. Subcategories define the clinical diagnosis. Cases with systemic AL amyloidosis were assigned to the AL-PCD category. This category included subcategories for AL amyloidosis diagnosed in the absence of other conditions (AL) or in the context of another hematologic condition such as multiple myeloma (AL/MM), chronic lymphocytic leukemia (AL/CLL), Waldenstrom’s macroglobulinemia (AL/WM) and light chain deposition disease (AL/LCDD). Diagnoses reported in the publications were not re-evaluated according to current diagnostic criteria. Cases lacking information about amyloidosis were assigned to the Other-PCD category, which included subcategories for MM, WM, LCDD, and POEMS (polyneuropathy, organomegaly, edema, M-protein and skin changes) syndrome. Twenty-nine sequences associated with asymptomatic or smoldering MM were grouped into the MM subcategory for the current analyses.

The third category in AL-Base is Non-PCD LCs, , which was originally intended to provide non-amyloidogenic control sequences [6]. These sequences were drawn from a diverse group of studies that exemplified the state of antibody sequencing technology in 2008, including studies of B cell malignancies such as CLL, the healthy immune response, and autoantibodies.The Non-PCD sequences, comprising 2,741 LCs, have been retained in AL-Base for reference. Six LC protein sequences associated with localized AL amyloidosis and one sequences from a case with hereditary AL amyloidosis were assigned to the Non-PCD category, since they were not derived from cases with a systemic PCD disorder.

### New monoclonal PCD LC sequences

PCD-associated LC sequences reported in the literature were identified by searching the NCBI PubMed, Nucleotide and Protein databases. Several sequences were identified that had not been included in the original version of AL-Base. Where possible, sequences were downloaded directly using the NCBI E-utilities system (Sayers 2022), implemented via the “rentrez” package in R (Winter 2017). Many sequences determined by Boston University have been updated to include the *IGC*_*L*_ region since their initial deposition. Sequences not deposited at NCBI were copied or manually transcribed from the publication. Additional LC sequences reported in proceedings of the International Symposia on Amyloidosis were manually curated. Nucleotide sequences were used in preference to protein sequences where both were available. Individual monoclonal LC sequences were named according to the convention of the institution where they were determined. All sequences were added to the AL-Base website.

Duplicate instances of the same clonal entity from the previous version of AL-Base were identified by sequence homology and removed. The longest and most complete sequence was retained, and nucleotide sequences were prioritized over protein sequences where possible. The earliest reference to each sequence was retained.

Several LC entries were deposited in NCBI as multiple identical or near-identical nucleotide sequences, which appear to represent either multiple colonies from a cloning procedure, or updated sequences where the original was not removed. These sequences were collapsed into a single consensus sequence, where possible, using multiple sequence alignments. Groups of sequences where an unambiguous consensus could not be identified were excluded. These consensus sequences were included in AL-Base with a note and list of the individual NCBI entities that were used to generate the consensus.

### Polyclonal LC sequences from Observed Antibody Space

Observed Antibody Space (OAS, https://opig.stats.ox.ac.uk/webapps/oas/) is a project that collects and annotates immune repertoires for use in large-scale analyses [26,27]. It contains over one billion sequences from over 80 different studies that are available for download based on specific attributes such as antibody chain type or disease state. Here, OAS was searched for polyclonal sequences to be used as a control group in analyses of PCD-associated LCs. Sixty-six experiments from six studies with >10,000 κ and λ LC protein sequences per experiment were identified. In each study, antibody cDNA libraries were created and sequenced using peripheral blood mononuclear cells from healthy or vaccinated individuals [29–34]. These data comprised 8,047,747 unique, non-redundant LC protein sequences covering complete *IGV*_*L*_-*IGJ*_*L*_ gene regions. In cases of redundant sequences, the longest was retained for analysis. This approach counted the number of distinct sequences derived from each *IGV*_*L*_ gene, rather than the number of clones within the immune repertoire, which cannot be directly measured. We assumed that these counts are similar, given the large number of individual sequences. The OAS sequences were not deposited in AL-Base.

### Alignment of LCs to germline precursor genes

The IMGT gene nomenclature and numbering was used throughout the manuscript [8]. For deposition in AL-Base,PCD-associated nucleotide sequences were aligned to the IMGT human germline database using the HighV-QUEST tool to identify the precursor germline genes and define the boundaries of the *IGV*_*L*_ and *IGJ*_*L*_ gene regions [35,36]. Protein sequences were aligned to the IMGT human germline database using the DomainGapAlign tool [37]. Only functional genes were used for germline assignment. Nucleotide sequences were also aligned using the NCBI IgBLAST tool [38], which gave concordant results in all cases. The *IGLJ2* and *IGLJ3* genes have indistinguishable sequences and cannot be unambiguously assigned from mRNA sequences. IgBLAST was used to determine the *IGC*_*L*_ precursor germline gene for nucleotide sequences, and DomainGapAlign was used to identify the constant(C_L_) domain for protein sequences. The *IGC*_*L*_ gene and C_L_-domain protein assignments were retained only when at least 150 nucleotides or 50 residues, respectively, were available. For the polyclonal control cohort, the *IGV*_*L*_ and *IGJ*_*L*_ gene assignments were retained as provided in OAS. Only *IGV*_*L*_*-IGJ*_*L*_ regions were analyzed from OAS sequences, since the original studies used sequencing strategies that do not provide complete *IGC*_*L*_ coverage.

### Analysis of LC precursor gene frequency

For the analyses presented here, each sequence was classified according to the extent of coverage of the *IGV*_*L*_ *-IGJ*_*L*_ region that encodes the V_L_-domain. We defined complete sequences as those with no missing or ambiguous amino acid residues. Incomplete sequences were defined as those missing N- or C-terminal residues, but which had unambiguous *IGV*_*L*_ and *IGJ*_*L*_ gene assignment and at least 80 contiguous residues, including coverage of all three complementarity determining regions (CDR). Sequences that did not meet these criteria were excluded from further analysis but retained in AL-Base for reference.

The individual LC sequences with both complete and incomplete *IGV*_*L*_*-IGJ*_*L*_ region coverage were counted for precursor germline gene analysis. Allele information was not considered. The *IGK* locus comprises two clusters, proximal and distal, and several genes have functional (i.e., non-pseudogene) paralogs in each cluster that cannot be distinguished by alignment of mRNA sequences [39]. To avoid bias from the alignment procedure, paralogous *IGKV* genes from the proximal and distal loci were considered identical and counted as derived from the proximal locus: e.g., *IGKV133* and *IGKV1D33* sequences were both counted as derived from the *IGKV133* gene.

The relative frequency of each *IGV*_*L*_ germline gene was calculated for AL, MM and OAS subcategories. The association of *IGV*_*L*_ germline gene with AL *vs*. MM or AL *vs*. OAS was tested by logistic regression analysis implemented via a generalized linear model in R v 4.2.2 [40]. Genes from both the *IGK* and *IGL* loci were considered together in this analysis. The odds ratio (OR) was used as a measure of the enrichment of each *IGV*_*L*_ gene for these comparisons and was considered an estimate for the amyloidogenic propensity of LC derived from that gene. Adjustment for multiple comparisons was performed using the Benjamini-Hochberg false discovery rate (FDR) method. FDR ≤ 0.05 was considered statistically significant. The correlation between gene frequencies in different subcategories was compared using the Pearson product moment correlation coefficient, *r;* a p-value ≤ 0.05 was considered statistically significant. Restricting the analysis to sequences with a complete *IGV*_*L*_*-IGJ*_*L*_ region did not alter the list of genes identified as significantly enriched.

### Mutational frequency analysis

The frequency of amino acid substitutions, insertions and deletions was calculated for each V_L_-domain with complete sequence coverage, relative to the translated sequence of its assigned *IGV*_*L*_*IGJ*_*L*_ alleles. The analysis considered amino acid changes, rather than nucleotide changes, so that LCs with only protein sequences could be included in the analysis. Therefore, the term “mutation” is used to refer to amino acid changes throughout the manuscript. We assumed that all differences between LC sequences and their assigned germline genes were due to somatic hypermutation, rather than undescribed variants in germline precursor genes.

Production of a functional LC protein is required for B cell survival [2], so mutations that introduce frameshifts or stop codons within the LC are not tolerated and have not been directly observed in AL amyloidosis. Mutation frequency was calculated as a fraction of the V_L_-domain length and expressed as a percentage:

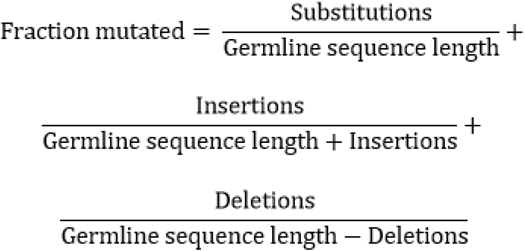

Mutation frequency was compared between AL, MM and OAS LC sequences, considering either locus (*IGL* or *IGK*) or germline *IGV*_*L*_ gene as a grouping variable. Non-parametric Kruskal-Wallace and Wilcoxon rank sum tests were used to assess the differences in mutation frequencies. Results were corrected for multiple comparisons, where FDR ≤ 0.05 was considered statistically significant.

### LC physicochemical properties

The isoelectric point and hydrophobicity of each complete LC sequence were calculated using the “peptides” R package [41]. For each parameter, the average value for all residues within the V_L_-domain was calculated. The default scales were used: EMBOSS PEPSTATS for pI and the Kyte-Doolitle algorithm for hydrophobicity. Non-parametric Kruskal-Wallace and Wilcoxon rank sum tests were used to assess the differences in parameters. Results were corrected for multiple comparisons, where FDR ≤ 0.05 was considered statistically significant.

### Selection of test cohort for amyloidogenicity predictions

To validate the performance of algorithms previously reported to predict LC amyloidogenic propensity [9–11], a new set of 1,590 V_L_-domain protein sequences (680 κ and 910 λ) with complete sequence coverage was compiled.

Three groups of sequences were used: (i) AL-Base AL sequences, which included AL and AL/MM subcategories (n = 155) (ii) AL-Base MM subcategory sequences (n = 784); and (iii) a random set of polyclonal sequences from OAS selected to match the number of sequences derived from each precursor gene in the AL subcategory from AL-Base, as of August 2023 (n=651). All PCD sequences were reported from 2020 to 2023 and were not tested in the original publications. Within this test cohort, AL sequences were defined as amyloidogenic, and MM and OAS sequences were defined as non-amyloidogenic.

### Performance of prediction algorithms

Sequences from the new validation dataset described above were submitted to the LICTOR [9], VLAmY-Pred [10] and AB-Amy [11] online servers. Since LICTOR did not accept κ or multiple LC sequence entries, an automated web query was used to submit λ sequences. Not all sequence submissions returned a prediction. Numbers of evaluated sequences are reported in Figure 5. The prediction from each algorithm was compared with the clinical subcategory. The sensitivity, specificity and accuracy were calculated and compared with parameters in the original publications:

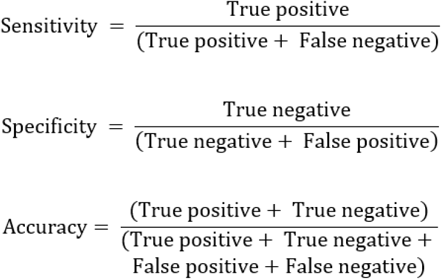

True positive and true negative represented correct identification of AL and non-AL sequences, respectively; false negative represented an unidentified AL sequence; and false positive represented a non-AL sequence identified as an AL sequence.

### Analysis software used

Most analysis was carried out using R v 4.2.2 [40] via the RStudio environment [42]. The following packages were used: Biostrings v 2.66.0 [43], broom v 1.0.3 [44], ggpubr v 0.6.0 [45], msa v 1.30.1 [46], rstatix v 0.7.2 [47], Peptides v 2.4.5 [41], polite v 0.1.3 [48], rentrez v 1.2.3 [49], rvest v 1.0.3 [50] and Tidyverse v 1.3.2 [51]D. For analysis of OAS sequences, the Unix Bash shell utilities *grep, sed* and *awk* were used on the Boston University Shared Computing Cluster.

## Results

### An updated AL-Base website and database

AL-Base contains LC nucleotide and protein sequences obtained from the literature and ongoing sequencing projects at the Boston University Amyloidosis Center. ALBase has been rebuilt to improve performance and provide new functionality. Each sequence has its own page with all associated data including an amino acid alignment with the assigned germline gene. Users can filter the database to generate a subset of sequences by clinical category or subcategory; *IGV*_*L*_ germline gene; κ or λ isotype; and the nucleotide or protein molecule type. The database can be searched for terms such as NCBI accession or specific sequence name. Alternatively, protein or nucleotide sequences within AL-Base can be searched via BLAST. Nucleotide or amino acid sequences can be downloaded individually from a sequence page or in bulk from the search results. The sequence can be directly exported to external tools including IgBLAST, Expasy ProtParam [52] and the IMGT tools V-QUEST, DomainGapAlign and Colliers-de-Perles [8]. From bulk search results, sequences can be aligned using Clustalw or IMGT alignment tools. The current number of sequences in AL-Base is graphically displayed in the Statistics section of the website.

Following addition of new and consolidation of existing sequences, there were 4,934 sequences in AL-Base as of August 2024. Of those, 2,193 were PCD-associated and 2,741 were Non-PCD-associated sequences. The distribution of sequences in different clinical categories and subcategories is shown in Table 1 and Figure 1. Complete *IGV*_*L*_*-IGJ*_*L*_ gene regions were identified in 1,788 (81.5%)PCD-associated sequences. A further 263 sequences(12%) had incomplete *IGV*_*L*_*IGJ*_*L*_ gene regions but could be assigned to a precursor gene, while the remaining 142 sequences (6.5%) had only partial sequences or non-productive *IGV*_*L*_*-IGJ*_*L*_ rearrangements. The latter sequences were excluded from further analyses, but retained in AL-Base for reference. The AL-PCD category comprised 847 complete or incomplete sequences associated with AL amyloidosis, and Other-PCD category was composed of 1,204 sequences from cases with plasma cell disorders without AL amyloidosis. Nucleotide sequences were available for 693 (82%) AL-PCD and 941 (78%) Other-PCD LCs. *IGLV* genes accounted for 73.3% of sequences in the AL-PCD category and 64.5% of sequences in the AL subcategory (Table 1 and Figure 1). The AL/MM subcategory had similar numbers of *IGKV* and *IGLV* sequences, representing 3.3% and 3.4% of the AL-PCD category, respectively. In the Other-PCD category, MM-associated sequences derived from *IGKV* genes accounted for 55.2% of sequences, while 37.9% were derived from *IGLV* genes. LCDD and WM sequences were predominantly from *IGKV* genes, and POEMS sequences were exclusively from *IGLV* genes.

**Table 1:** LC sequences in AL-Base. Categories define the clonal origin of the LC and whether amyloidosis diagnosis was reported. Subcategories define individual diseases. “AL” represents AL amyloidosis as an independent diagnosis or in the context of another underlying malignancy. MM, multiple myeloma; CLL, chronic lymphocytic leukemia; LCDD, light chain deposition disease; POEMS, polyneuropathy, organomegaly, edema, M-protein and skin changes syndrome; WM, Waldenström macroglobulinemia. “Other” includes Fanconi syndrome without overt MM and the LC sequence from a clone associated with heavy chain deposition disease. “Localized AL” represents localized AL amyloidosis, which is not associated with a PCD. Other Non-PCD sequences include polyclonal sequences from healthy individuals and those with infectious or autoimmune diseases; and clonal sequences from other B cell malignancies. Incomplete *IGV*_*L*_*-IGJ*_*L*_ region coverage is defined as having at least 80 contiguous residues, including with complete coverage of CDRs 1-3.

| Category | Subcategory | Complete $IGV_L$-$IGJ_L$ region | | | Incomplete $IGV_L$-$IGJ_L$ region | | | Fragments or non-productive genes | | | Total |
| --- | --- | --- | --- | --- | --- | --- | --- | --- | --- | --- | --- |
| | | $IGKV$ | $IGLV$ | Total | $IGKV$ | $IGLV$ | Total | $IGKV$ | $IGLV$ | Total | |
| AL-PCD<br>n = 891 | AL | 160 | 524 | **684** | 34 | 63 | **97** | 6 | 35 | **41** | **822** |
|  | AL/MM | 29 | 25 | **54** | 0 | 4 | **4** | 0 | 2 | **2** | **60** |
|  | AL/CLL | 0 | 2 | **2** | 0 | 0 | **0** | 0 | 0 | **0** | **2** |
|  | AL/LCDD | 1 | 0 | **1** | 0 | 0 | **0** | 0 | 0 | **0** | **1** |
|  | AL/WM | 2 | 3 | **5** | 0 | 0 | **0** | 1 | 0 | **1** | **6** |
|  | Total | 192 | 554 | **746** | 34 | 67 | **101** | 7 | 37 | **44** | **891** |
| Other-PCD<br>n = 1302 | MM | 595 | 374 | **969** | 70 | 83 | **153** | 42 | 26 | **68** | **1190** |
|  | LCDD | 29 | 4 | **33** | 3 | 0 | **3** | 2 | 4 | **6** | **42** |
|  | POEMS | 0 | 31 | **31** | 0 | 5 | **5** | 0 | 13 | **13** | **49** |
|  | WM | 5 | 0 | **5** | 0 | 0 | **0** | 8 | 0 | **8** | **13** |
|  | WM/LCDD | 0 | 0 | **0** | 1 | 0 | **1** | 0 | 0 | **0** | **1** |
|  | Other | 4 | 0 | **4** | 0 | 0 | **0** | 3 | 0 | **3** | **7** |
|  | Total | 633 | 409 | **1042** | 74 | 88 | **162** | 55 | 43 | **98** | **1302** |
| Non-PCD<br>n = 2741 | Localized AL | 0 | 1 | **1** | 1 | 2 | **3** | 0 | 2 | **2** | **6** |
|  | Non-PCD | 1051 | 581 | **1632** | 621 | 346 | **967** | 102 | 34 | **136** | **2735** |
|  | Total | 1051 | 582 | **1633** | 622 | 348 | **970** | 102 | 36 | **138** | **2741** |

**Figure 1:**
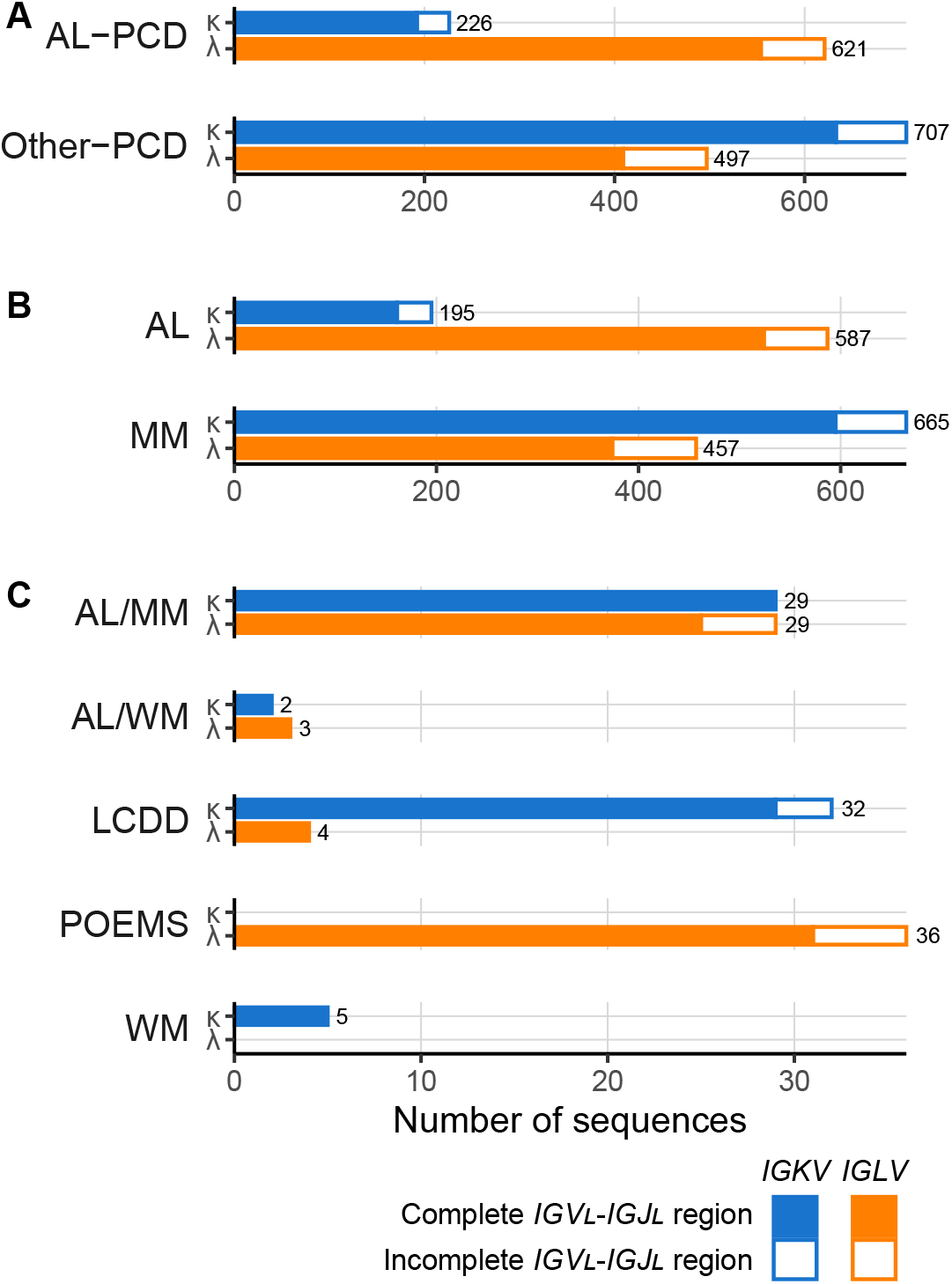
PCD-associated LC sequences in AL-Base, according to clinical categories (A) and disease subcategories (B and C), defined as for Table 1. Data for κ LCs are shown in blue and data for λ LCs are shown in orange. Solid bars represent sequences with complete *IGV*_*L*_*-IGJ*_*L*_ gene region coverage. Hollow bars represent sequences with unambiguous *IGV*_*L*_ and *IGJ*_*L*_ gene assignment but incomplete *IGV*_*L*_*-IGJ*_*L*_ gene region coverage. Total sequence numbers are shown for each bar.

The Non-PCD category comprised 2,741 sequences, which were included in the original AL-Base as non-amyloidogenic controls, and six LC sequences associated with localized AL amyloidosis. For the current analyses, polyclonal sequences from OAS were used instead of those from the Non-PCD category. The Non-PCD sequences have been retained in AL-Base for reference. From the OAS database we identified 8,047,747 unique *IGV*_*L*_*-IGJ*_*L*_ sequences, 4,278,425 *IGKV-IGJV* (53%) and 3,769,322*IGLV-IGJV* (47%), derived from healthy peripheral B cells, which are referred to as “OAS” in the analyses below.

### A subset of IGV_L_ precursor germline genes is more frequent in AL compared to MM or the polyclonal OAS repertoire

The association between precursor IGVL genes and amyloidosis was assessed by counting the number of LCs in the AL subcategory derived from each gene, and measuring their enrichment relative to the MM subcategory or the OAS repertoire. Precursor gene frequency varied among PCD clones and the polyclonal repertoire (Figures 2 and 3). Within the AL subcategory, 482 of 781 sequences (61.6%) were derived from five precursor germline genes, *IGKV1-33, IGLV1-44, IGLV2-14, IGLV3-1* and *IGLV6-57*. These five genes accounted for 316 of 1,122 MM sequences (28.2%) and 1,424,811 of 8,047,747 OAS sequences(17.8%).

**Figure 2:**
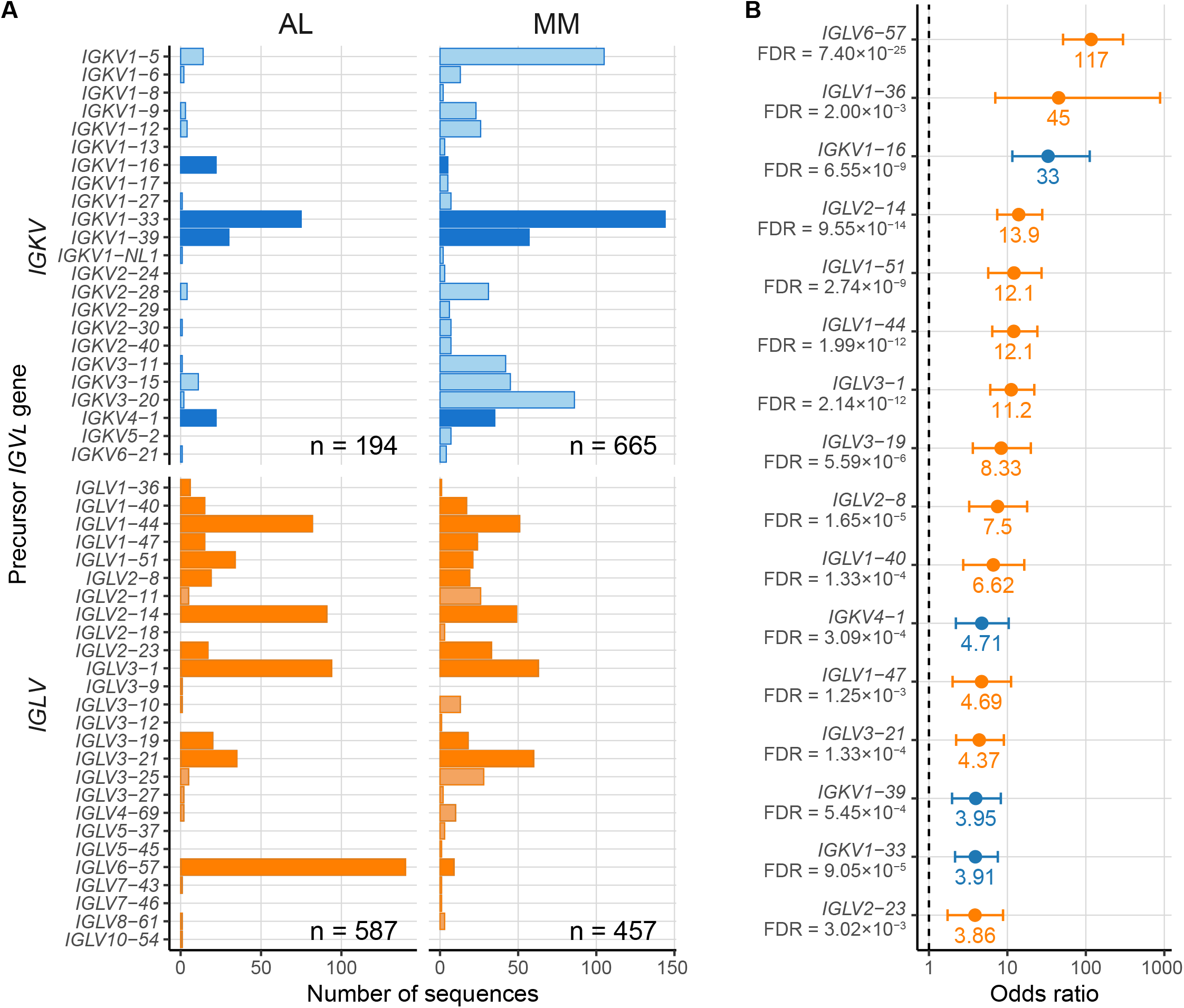
*IGV*_*L*_ precursor germline gene use differs between AL- and MM-associated sequences. Data for *IGKV* and *IGLV* genes are shown in blue and orange, respectively. A) Numbers of individual *IGV*_*L*_ sequences derived from each precursor gene. Dark colored bars represent *IGV*_*L*_ genes significantly over-represented in AL *vs*. MM sequences. B) Odds ratios for *IGV*_*L*_ precursor germline genes significantly over-represented in AL *vs*. MM sequences. Error bars show 95% confidence intervals. FDR, false discovery rate; FDR ≤ 0.05 was considered statistically significant.

**Figure 3:**
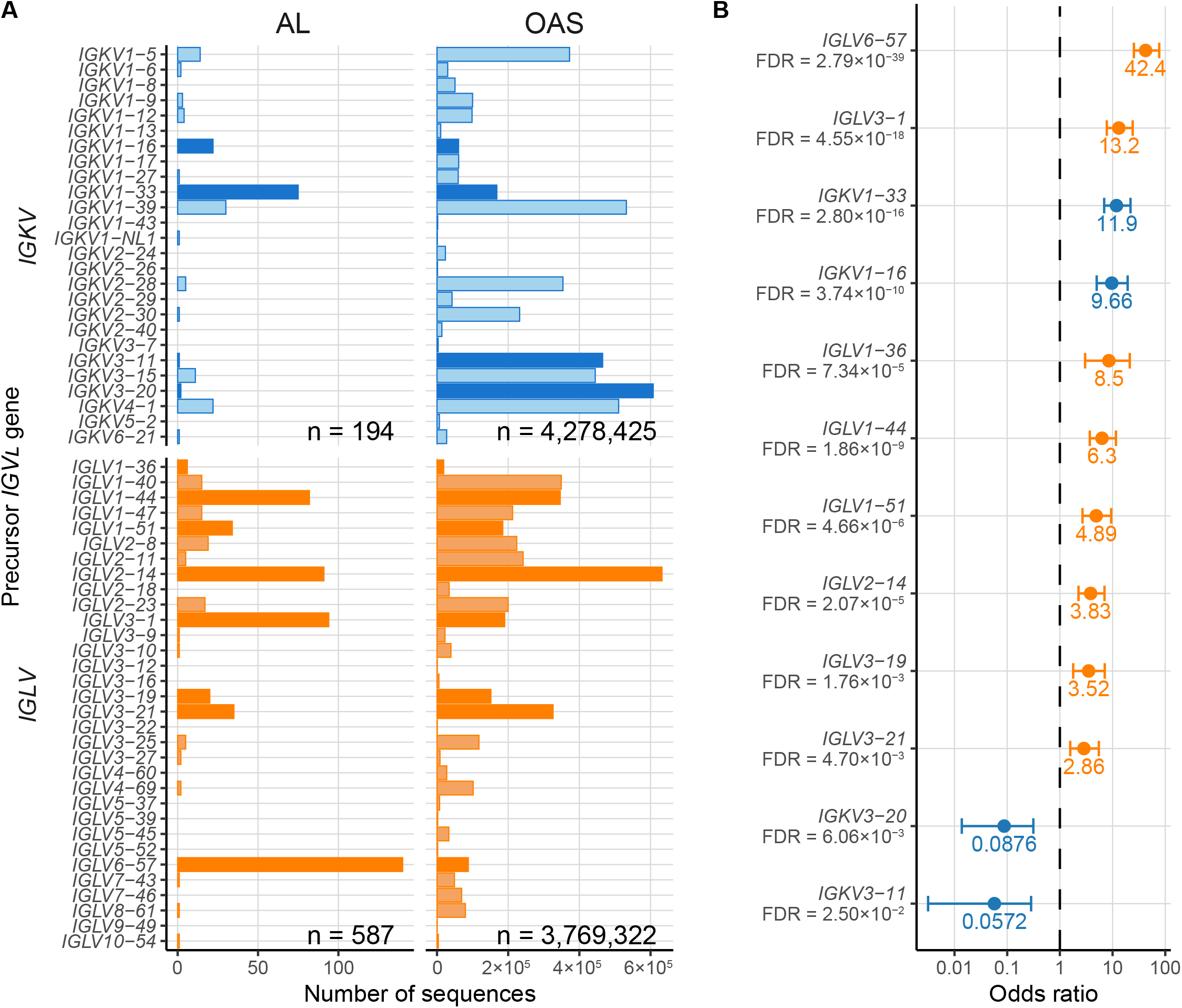
*IGV*_*L*_ precursor germline gene use differs between monoclonal AL- and polyclonal OAS-associated sequences. Data for *IGKV* and *IGLV* genes are shown in blue and orange, respectively. A) Numbers of individual *IGV*_*L*_ sequences derived from each precursor gene. Dark colored bars represent *IGV*_*L*_ genes significantly over-represented in AL *vs*. OAS sequences. B) Odds ratios for *IGV*_*L*_ precursor germline genes significantly over- or under-represented in AL *vs*. OAS sequences. Error bars show 95% confidence intervals. FDR ≤ 0.05 was considered statistically significant.

The comparison of precursor *IGV*_*L*_ gene use in the AL *vs*. MM subcategories is shown in Figure 2. The overall pattern of gene use was correlated between these two groups (*r* = 0.48; p = 0.00015). Sixteen *IGV*_*L*_ genes were significantly enriched in AL *vs*. MM sequences after adjustment for multiple comparisons. The strongest association with AL amyloidosis was observed for *IGLV6-57* (OR 117; 95% CI 51.2-298), followed by *IGLV1-36* (OR 45; 95% CI7.01-884) and *IGKV1-16* (OR 33; 95% CI 11.6-112) (Figure 2B). Other statistically significant associations included ten *IGLV* (*IGLV1-40, IGLV1-47, IGLV1-51, IGLV1-44, IGL2-08, IGLV2-14, IGLV2-23, IGLV3-01, IGLV3-19*,and *IGLV3-21*) and three *IGKV* (*IGKV1-33, IGKV1-39*and *IGK4-01*) germline genes with OR ranging from 13.9to 3.86. No genes were significantly under-represented in the AL *vs*. MM subcategories. The frequency of *IGJ*_*L*_ and *IGC*_*L*_ gene use was similar between the AL and MM subcategories.

The 58 AL/MM sequences were initially excluded from the analysis. The fraction of AL/MM and AL LCs derived from each precursor gene was similar (*r* = 0.74; p = 2.1×10^-11^). An exception was *IGLV657*, which accounted for only 3 of 58 (5%) AL/MM sequences. Combining the AL and AL/ MM subcategories (AL + AL/MM) did not alter the list of over-represented genes compared to MM.

MM-associated LC sequences may be biased by the proliferative nature of the MM clone. Furthermore, several genes are infrequent in MM clones, leading to large uncertainties in the calculated odds ratios. To investigate whether biases in the MM sequences influence the apparent over-representation of germline genes in AL, the fraction of AL LC sequences derived from each germline gene was compared with polyclonal OAS sequences. The correlation coefficient between AL and OAS was 0.37 (p = 0.0039), indicating that gene use in AL is less similar to OAS than to MM. Eight *IGLV* (*IGLV1-36, IGLV1-44, IGL1-51, IGLV2-14, IGLV3-01, IGLV3-19, IGLV3-21*, and *IGLV6-57*) and two *IGKV* (*IGKV1-16* and *IGKV1-33*) genes were significantly over-represented, and two *IGK* genes (*IGKV3-11* and *IGKV3-20*) were significantly under-represented in the AL *vs*. OAS comparison (Figure 3).

### Somatic mutation frequency differs between AL, MM and the polyclonal repertoire

Antibodies expressed by plasma cells have undergone somatic hypermutation in response to antigen, accumulating multiple amino acid changes relative to their germline regions [2]. Amyloidogenic LCs have been hypothesized to harbor more mutations than non-amyloidogenic sequences [9,53,54]. To test this hypothesis, the frequency of residue replacements, insertions and deletions, corrected for sequence length, was calculated for AL, MM and OAS LCs with complete V_L_-domain sequence coverage. LCs had a wide range of mutation frequencies (Figure 4). Among AL-associated LCs, the frequency of mutations varied from 2.7% to 35% for λ V_L_-domains and 1.9% to 22% for κ V_L_-domains. For MM-associated LCs, ranges were 1.8% to 27% for λ V_L_-domains and 0% to 25% for κ V_L_-domains. The OAS repertoire contains a small fraction of highly mutated sequences, including sequences with large insertions, which extends the apparent range of mutation frequencies. However, 95% of the sequences have mutation frequencies of 1.9–21% for κ and 1.8-24% for λ V_L_-domains.

**Figure 4:**
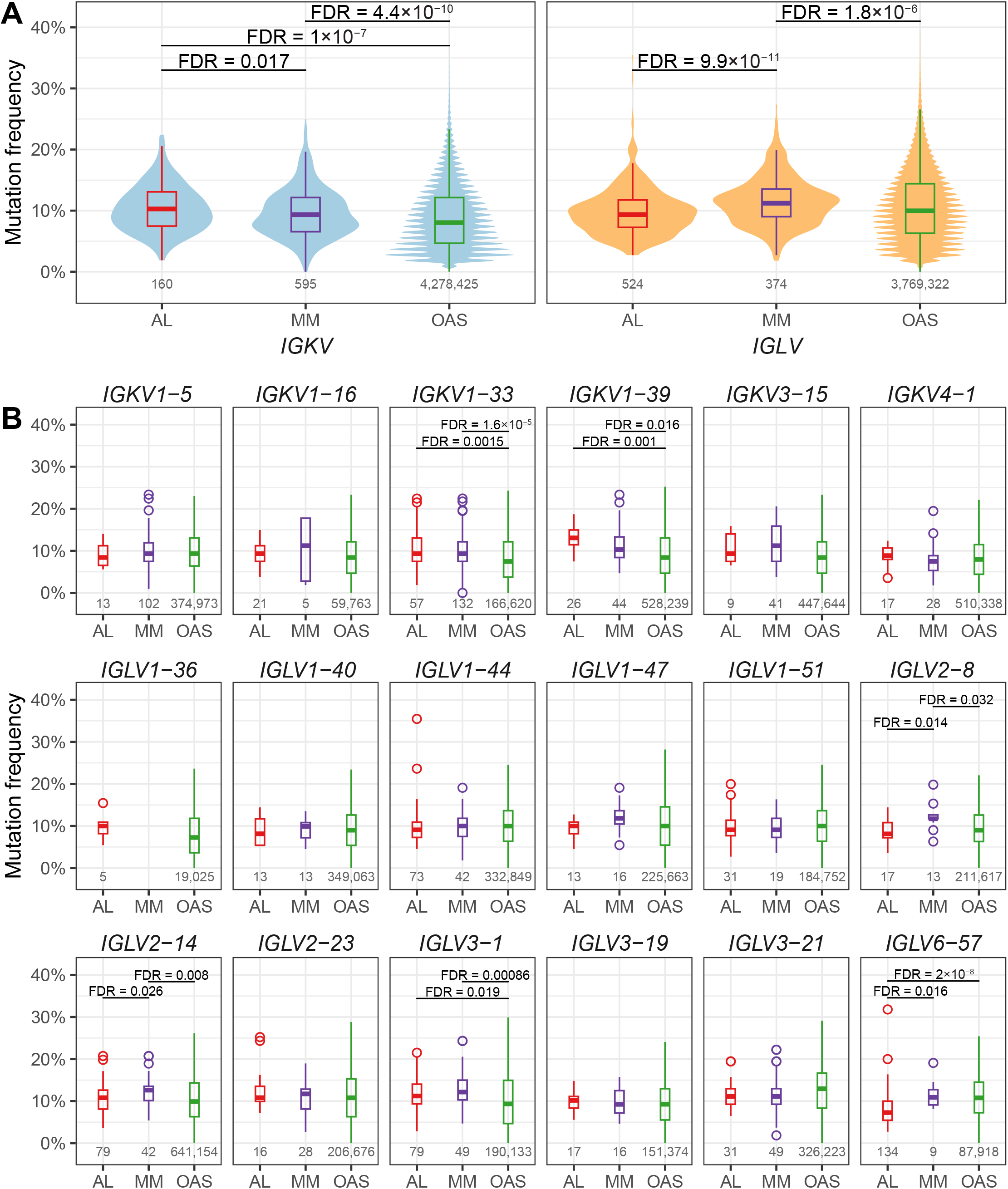
Mutation frequency varies between LCs associated with AL, MM and OAS. The number of amino acid residue replacements, insertions and deletions for each V_L_-domain with complete sequence coverage is expressed as a percentage of length. The number of sequences in each group is shown below the boxplots. Significant differences are shown after adjustment for multiple comparisons (FDR ≤ 0.05). A) Comparison for κ and λ LC sequences. Violin plots show the range and distribution of mutation frequency. B) Comparison for LCs derived from 18 *IGV*_*L*_ germline genes with at least five AL-associated sequences. The box and whisker plots show median (central bars), inter-quartile range (boxes) and distance to the non-outlier data (whiskers). Circles show outliers for AL and MM sequences.

Upon stratification by *IGV*_*L*_ locus, significant differences in mutational frequency were observed between the AL, MM and OAS groups after correction for multiple testing (Figure 4A). The median mutation frequencies were 10.3%, 9.4%, 8.0% for κ and 9.4%, 11.2% and 10.0% for λ V_L_-do-mains in the AL, MM and OAS subcategories, respectively. Among κ V_L_-domains, a significantly higher mutation frequency was found in AL *vs*. MM and AL *vs*. OAS sequences, whereas among λ V_L_-domains, the mutation rate was significantly lower in AL *vs*. MM sequences.

A similar pattern was observed for mutation frequencies between individual germline genes after correction for multiple testing (Figure 4B). Sequences derived from *IGKV1-33* and *IGKV1-39* had a significantly higher mutation rates in the AL *vs*. OAS comparison, with median differences of 2.8% and 3.7%, respectively. Sequences derived from *IGLV2-8, IGLV2-14* and *IGLV6-57* had significantly lower mutation frequencies in the AL *vs*. MM comparison, with median differences of 1.8%, 3.6% and 3.6%, respectively. Sequences derived from *IGLV3-1* and *IGLV6-57* had a significantly lower mutation rates in AL *vs*. OAS subcategories, with median differences of 1.9% and 2.7%, respectively.

### Physicochemical properties are similar between AL, MM and the polyclonal OAS repertoires

Physicochemical properties have been proposed to contribute to amyloidogenic potential of monoclonal LCs [3]. To investigate these effects, the theoretical isoelectric point and hydrophobicity score were calculated for AL and MM sequences with a complete V_L_-domain coverage. Although variations in these parameters were found between LC sequences derived from different *IGV*_*L*_ genes, no significant differences were identified in AL *vs*. MM subcategories within each gene after correction for multiple testing.

### Amyloidogenicity prediction tools demonstrate reduced accuracy on new sequences

Machine learning algorithms have been reported to predict the amyloidogenic propensity of LC sequences [9–11]. The performance of these algorithms was tested with a new dataset of 1590 AL, MM and polyclonal OAS V_L_-domain sequences. The sequences were submitted to freely-available web-based prediction servers that use various machine learning approaches: VL-AmyPred [10] and Ab-Amy [11] to evaluate both κ and λ sequences, and LICTOR [9] forλ sequences. The performance of these algorithms on the new dataset was compared to that previously described in the publications (Figure 5). The accuracy, sensitivity and specificity of each algorithm was lower for the new dataset compared to dataset that was initially used to train the algorithms.

**Figure 5:**
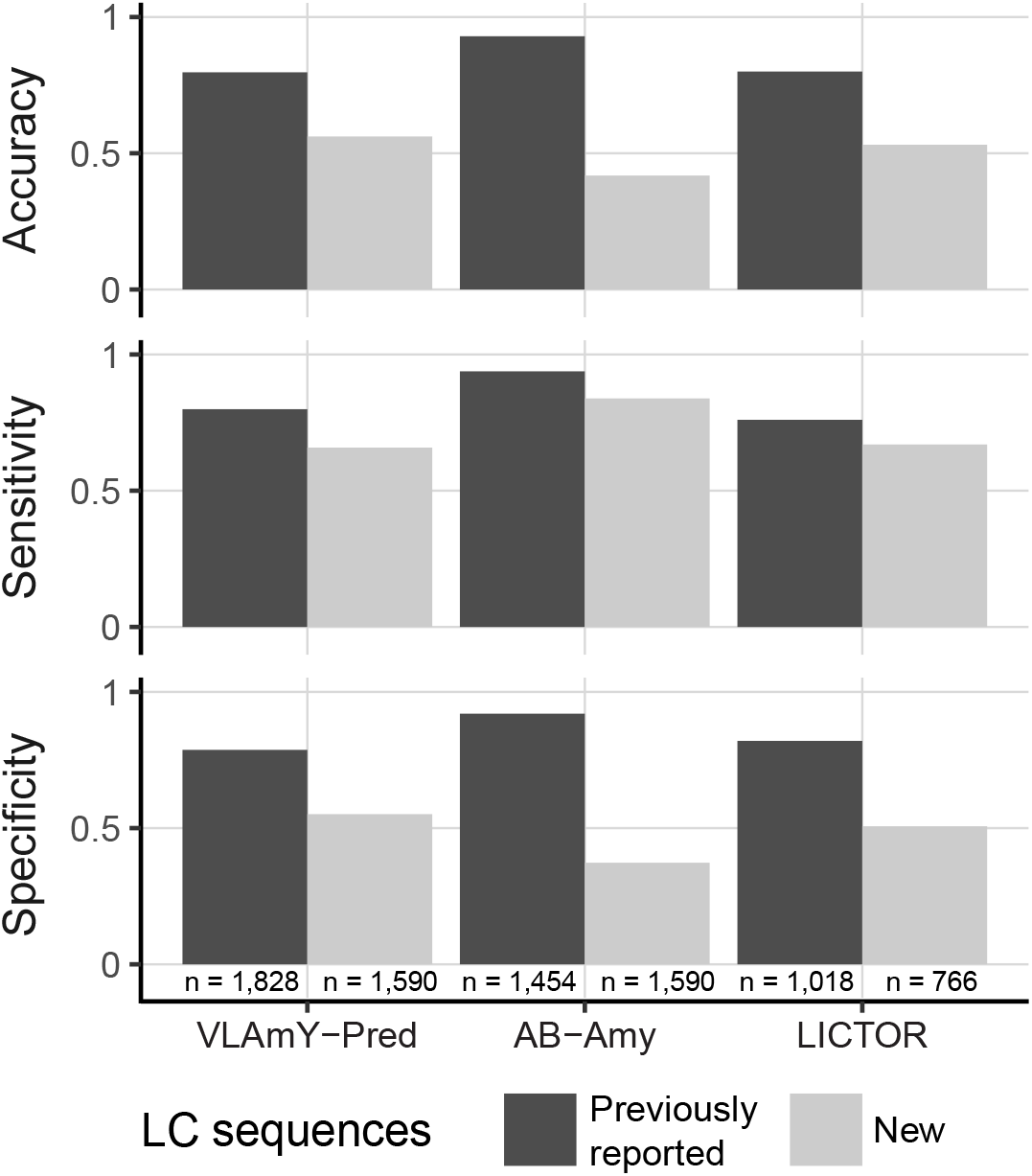
Amyloidogenicity prediction tools achieve lower accuracy, sensitivity and specificity with a new dataset of AL, MM and polyclonal OAS V_L_-domain sequences. The parameters reported for each algorithm with the original sequences are shown with dark bars, and the parameters calculated for 1,590 new sequences are shown withlight bars. Note that LICTOR algorithm accepts only λ sequences.

## Discussion

We present an updated AL-Base, a database of monoclonal LC sequences to study PCDs including AL amyloidosis. Of 2,193 PCD-associated LC sequences, 1,788 have complete nucleotide or amino acid coverage of *IGV*_*L*_*-IGJ*_*L*_ regions. A total of 847 AL- and 1,122 MM-associated sequences allows exploration of the connections between LC features and AL amyloidosis. As an additional control, 8,047,747 polyclonal OAS sequences were identified, facilitating additional analyses. A subset of sixteen LC precursor germline genes was enriched in AL clones, relative to MM clones (Figure 2) Most of these genes were also prevalent in AL when compared to the polyclonal OAS repertoire (Figure 3). Mutation frequency varied widely between LCs (Figure 4). However, among κ V_L_-domains, AL-associated LCs had significantly more mutations than MM-associated LCs, whereas the opposite was observed for λ V_L_-domains. Tools that predict amyloidogenicity performed less well on a new set of sequences than on the date that was originally used in their development (Figure 5).

AL amyloidosis is invariably preceded by an asymptomatic precursor phase known as monoclonal gammopathy of uncertain significance (MGUS) [55,56]. Incidence of AL amyloidosis can reach 1% in MGUS cases and may be higher among individuals with asymptomatic (smoldering) MM [1]. Therefore, there is a window of opportunity for early diagnosis and intervention to prevent amyloid deposition and organ damage. The association of a subset of *IGV*_*L*_ genes with amyloidosis means that LC sequencing is a promising approach for identifying individuals at risk for AL amyloidosis. A study designed to sequence the *IGLV* precursor genes in cases with MGUS and smoldering MM, has identified one AL amyloidosis case derived from the *IGLV2-14* precursor out of 17 studied patients [57]. To our knowledge, no other studies of *IGV*_*L*_ gene use in MGUS are currently available and, therefore, the frequency of genes representing the risk for AL amyloidosis in this cohort is unknown.

Several studies have investigated the association of *IGV*_*L*_ gene use with AL amyloidosis [6,9–16,58]. Similarly to previous reports, the *IGKV1-33, IGLV1-44, IGLV2-14, IGLV3-1* and *IGLV6-57* germline genes accounted for most AL clones in current analysis (Figures 2 and 3). The frequency of each *IGV*_*L*_ precursor gene in the AL-Base repository is highly correlated with the frequencies observed in the mass spectrometry analysis of amyloid deposits (*r*= 0.97, p = 9.6×10^9^) [12]. This data supports the hypothesis that a subset of *IGV*_*L*_ precursor genes is consistently over-represented in AL amyloidosis, though the studies may not represent completely independent populations as a subset of individuals was likely studied by both methods.

The non-AL-associated sequences in this report allowed quantification of *IGV*_*L*_ gene enrichment in AL amyloidosis compared to monoclonal MM-associated or polyclonal OAS sequences. We hypothesize that the ORs calculated for each *IGV*_*L*_ gene in Figures 2B and 3B represent a measure of relative risk for AL amyloidosis in the absence of other factors. The *IGLV6-57, IGKV1-16* and *IGLV1-36* genes that are most highly enriched in AL amyloidosis are relatively rare in MM and the OAS repertoire, suggesting that their expression may represent a risk factor for AL amyloidosis in individuals with MM or MGUS. An important caveat is that the rarity of LCs derived from these genes in the MM subcategory means that the uncertainty associated with their enrichment is high. The observation that these genes are also significantly enriched in AL *vs*.

OAS supports the hypothesis that they represent a risk for amyloidosis. A proteomics study identified nine cases of systemic AL amyloidosis associated with *IGKV1-16*, eight of which had cardiac involvement [12]. This study did not report the number of LCs derived from *IGLV1-36*, grouping this gene together with other rare *IGVL* genes [12]. LC sequences derived from the *IGKV3-11* and *IGKV3-20* precursor genes were significantly under-represented in AL *vs*. OAS, consistent with a low amyloidogenicity for these genes, which are among the most frequently observed in the OAS repertoire. Additionally, this analysis showed that some *IGV*_*L*_ genes, notably *IGKV1-33* and *IGLV2-14*, were frequently observed in both AL and MM clones, highlighting that the identity of precursor gene alone is not sufficient to determine the risk of AL amyloidosis.

Studies of multiple LC proteins have hypothesized that AL-associated sequences are more frequently mutated than non-AL-associated sequences [9,53,54]. The monoclonal LC sequences expressed by plasma cells are expected to undergo somatic hypermutation and acquired a larger number of mutations than sequences expressed by less differentiated B cells [2]. Therefore, the comparison between AL- and MM-associated sequences, which are both derived from bone marrow plasma cells, may be more meaningful than that between AL-associated sequences and all available LCs. The number of MM-associated LC sequences currently available in AL-Base allows a robust analysis of mutation frequency. This analysis revealed that the overall frequency of residue changes varied widely between sequences, but V_L_-domains derived from the *IGKV* genes have significantly more mutations, and V_L_-domains derived from the *IGLV* genes have significantly less mutations in the comparison between AL and MM sequences (Figure 4). Similar trends showed V_L_-domain sequences derived from a number of individual *IGKV* (*IGKV1-33* and *IGKV1-39)* and *IGLV (IGLV2-8, IGLV2-14, IGLV3-1 andIGLV6-57)* genes (Figure 4). LCs derived from *IGLV6-57* have a low mutation frequency, consistent with the reported high amyloidogenic propensity of V_L_-domain sequences encoded by *IGLV6-57* germline precursor [59–61]. Overall, AL-associated V_L_-domain sequences demonstrated no major differences in the frequency of somatic mutations compared to MM sequences, consistent with the previous observations that the position and character of residue changes are more likely to contribute to LC amyloidogenicity than their absolute numbers [9,62]. Polyclonal OAS sequences showed a wider distribution of mutation frequencies than PCD-associated sequences (Figures 4A), and the original dataset from the Non-PCD category was enriched in sequences with fewer mutations. This is consistent with low mutational load in sequences derived from the heterogeneous populations of B cells from which both control cohorts were derived [6,29–34]. Although the Non-PCD sequences remain available in the AL-Base repository, we recommend that future studies use large-scale repertoire sequencing data that are available from OAS resource.

In line with previous reports, the analysis of pI and hydrophobicity score values revealed no difference betweenV_L_-domain sequences from AL and MM subcategories [24]. Although physicochemical properties determine the ability of LCs to aggregate [3], these differences are not captured by the sequence-based metrics that were considered here.

Prediction of amyloid propensity requires testing a wide range of LC sequences (Figure 5). The apparent performance of predictive tools is strongly dependent on the proportion of AL- and non-AL-associated sequences in the test set [63]. The previously reported prediction algorithms were trained on the dataset of LC sequences from the old version of AL-Base and therapeutic antibody sequences, which captured only a fraction of the potential LC sequence diversity [9–11]. A large number of non-AL LC sequences was deliberately added to our test set to validate prediction algorithms. Our analysis did not confirm the original performance metrics of VL-AmyPred, Ab-Amy or LICTOR tools to reliably distinguish between AL- and non-AL-associated sequences (Figure 5) [9–11]. The performance of each algorithm was lower with the new sequences, as has been observed with other algorithms that are over-trained on their initial data [64]. The expanded AL-Base data presented here provides a resource for training the next generation of amyloid prediction tools and creation of new models. As more AL-associated sequences become available in the future, the performance of these tools will continue to improve. We anticipate that a successful diagnostic tool may include clinical information along with LC sequence data.

This study has several limitations: (i) At a fundamental level, the sample size of available PCD sequences remains relatively small compared to LC diversity, which was demonstrated by the broad range of confidence intervals in a subset of AL-associated rare germline gene precursors. (ii) The classification of sequences depends on the clinical information reported at the time of publication. The definition of PCD disorders has changed since older sequences were determined and patients were first diagnosed [65]. Hence, a proportion of sequences previously classified as AL may be considered AL/MM by current diagnostic criteria, or vice-versa. Importantly, the amyloid deposition that is relatively common in MM may not be looked for, detected or reported [66]. AL amyloidosis has been reported in 10-15% and amyloid deposits have been identified in 38% of patients with MM [4,67]. Thus, the assignment of LC sequence into the Other-PCD category only indicates the lack of diagnosis of AL amyloidosis, but does not exclude the presence of amyloid deposits. (iii) The amino acid sequences of only V_L_-domain were considered for our analyses, while the C_L_-domains and, perhaps, heavy chain partners may also contribute to amyloid propensity [68–70]. (iv) The OAS sequences were derived from a population of peripheral blood B cells, rather than bone marrow plasma cells, which may alter the distributions of germline gene usage and mutation frequency. (v) The results of statistical tests should be interpreted with the caveat that the AL-Base sequences are not representing a truly random set of monoclonal LC proteins, since sequences were compiled after the diagnosis from multiple studies with a variety of selection criteria.

Looking ahead, it is likely that the number of PCD-associated LC sequences will increase in coming years as high throughput sequencing of clonal genes, characterization of plasma cells by RNA sequencing and characterization of monoclonal proteins by mass spectrometry become more widespread [19–21]. We will continue to update AL-Base as more sequences become available. The set of LC sequences analyzed in current report can be used in other studies with the expectation that new, independent data will become available for validation. To this end, we invite investigators involved in AL amyloidosis research to deposit LC sequences into public repositories from where they that can be retrieved and incorporated into AL-Base for convenient access by the wider scientific community.

## Acknowledgements

This work was made possible by the generosity of patients who allowed their cells and sequences to be used for research, and scientists who shared LC sequences in publications and repositories. We thank Drs. Paola Rognoni, Francesca Lavatelli and Mario Nuvolone from the Amyloidosis Research and Treatment Center at University of Pavia, Italy; Dr. Raymond Comenzo from the Tufts University School of Medicine, USA; Dr. Marina Ramirez-Alvarado from the Mayo Clinic, USA; Dr. Stefan Schönland from the Amyloidosis Center at Heidelberg University Hospital, Germany; and Drs. Christophe Sirac and Vincent Javaugue from the Université de Limoges, France, for helping to collect and compile sequences generated from their institutions. The AL-Base website was built and is currently managed by the Biostatistics and Epidemiology Data Analytics Center at the Boston University School of Public Health.

